# SH2*scan*: Mapping SH2 domain-ligand binding selectivity for inhibitors and degraders

**DOI:** 10.1101/2025.07.09.663991

**Authors:** Luis M. Gonzalez Lira, Jennifer K. Wolfe-Demarco, Alexander M. Clifford, Tuan D. Le, Ghadeer M. Khasawneh, Medhanie Kidane, Michelle Nguyen, Julia A. Najera, Gabriel Pallares, Nicole B. Servant, Jean A. Bernatchez

## Abstract

Drug discovery targeting SH2 domains (key protein-protein interaction modules) has been hampered by a lack of assay systems evaluating synthetic ligand binding selectivity toward SH2 domains, to reduce potential off-target effects. In addition, the molecular determinants for synthetic ligand engagement to SH2 domains across the target class have yet to be defined. Here, we developed SH2*scan*, a high-throughput competition binding assay platform to quantify ligand-SH2 domain interactions, covering >80% of the target class. We uncovered unique binding selectivity profiles and quantified a broad range of dissociation constants (KDs) for 9 synthetic ligands of SH2 domains from the scientific literature with a range of reported primary targets. These results demonstrate that SH2*scan* can be used to design more selective compounds targeting SH2 domains. The platform can be further leveraged for the discovery of new molecular probes for the dissection of cellular protein-protein interaction networks.

## Main

Protein-protein interactions (PPIs) have reemerged as prime targets for drug discovery programs, given their importance as key biochemical transducers in deregulated cell signaling pathways, as is the case in inflammatory disorders and cancer^1,2^. SH2 domains, of which there are 120 canonical members found in 110 proteins, are prototypical PPI modules which generally recognize phosphotyrosine-containing motifs on their protein binding partners^3–5^. Despite divergent amino acid sequences among SH2 domains, the conserved structure among these PPI modules has presented challenges in the development of highly selective inhibitors of targets in the class^6^. While protein microarrays evaluating SH2 domain-phosphopeptide ligand interactions have been reported^7,8^, large- scale platforms to study the selectivity of SH2 domain interactions with small molecule inhibitors or targeted protein degraders have yet to be developed. As such, the molecular determinants for synthetic ligand binding selectivity across the SH2 domain target class remain largely unknown. Systems that would allow for the study of these features would be of high value to drug discovery programs targeting SH2 domains, given their potential to identify off-target liabilities for hit-to-lead programs in the early stages of drug development. Furthermore, many SH2 domain-containing proteins have limited biochemical and functional characterization in the scientific literature. Platforms which can identify selective small-molecule probes that can be subsequently used to interrogate the function of SH2 domain-containing gene products would be of significant value to the chemical biology community.

Lastly, modular and on-demand screening of phosphopeptide ligands derived from cellular proteins against SH2 domains at scale would be useful in the delineation of new PPI networks within the cell and could be used to gain new biological insight into signaling networks for therapeutic intervention.

Here, we describe a high-throughput, plate-based competition binding assay platform called SH2*scan*, with coverage of over >80% of the canonical human SH2 domains. We report the selectivity binding profiles of 9 synthetic ligands (8 SH2 domain binders and 1 proteolysis targeting chimera (PROTAC) which engages an SH2 domain on its target) (**Fig. 1**) from the literature in our assay platform. The data reveal unique binding selectivity signatures and a range of dissociation constants (KDs) across SH2 domains for these compounds. These results demonstrate the utility of the SH2*scan* platform in delineating the chemical determinants of SH2 domain binding selectivity for synthetic ligands across the target class.

**Fig. 1.**
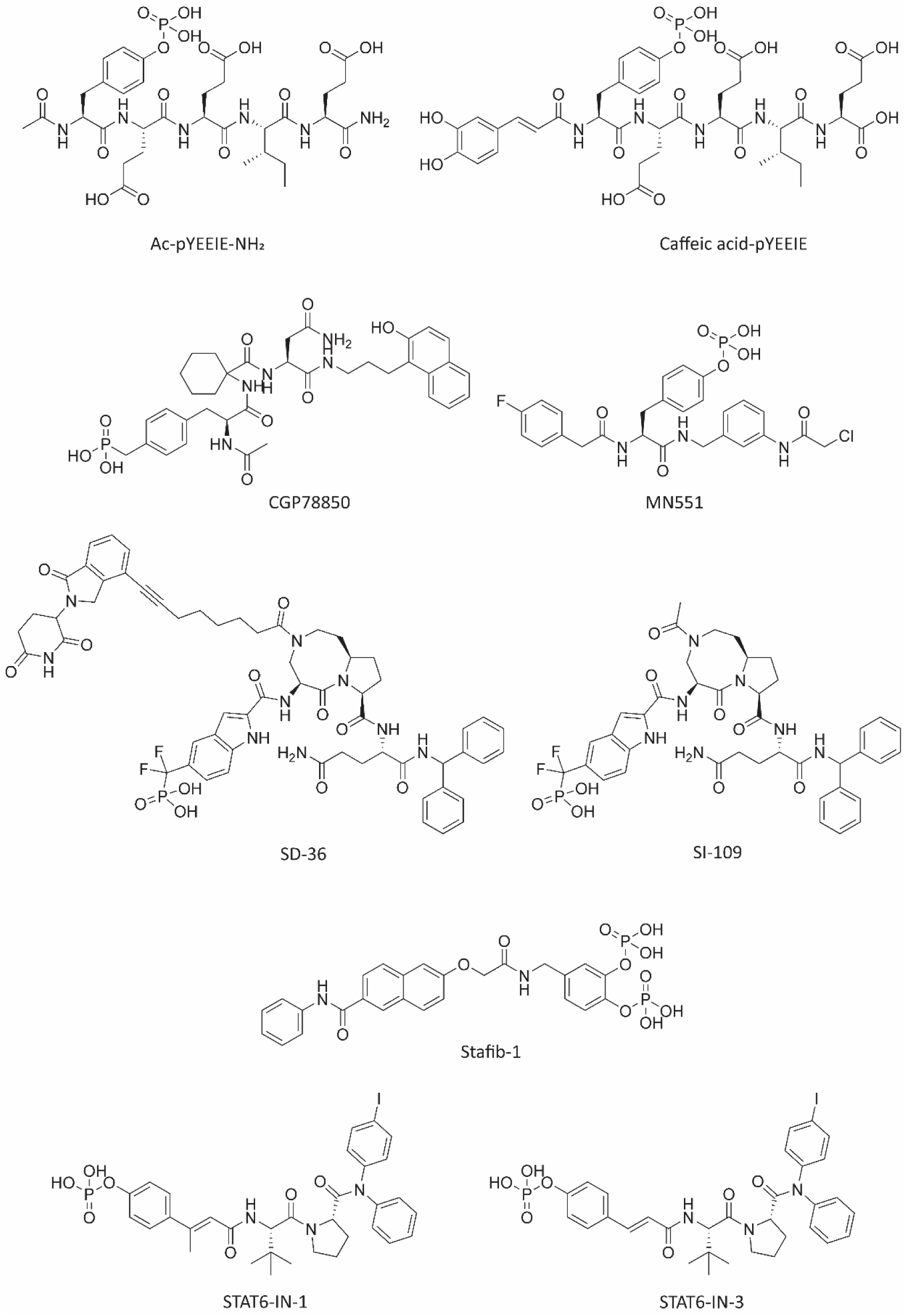
Chemical structures of the 9 synthetic SH2 domain ligands tested in SH2*scan*. Compounds tested for binding selectivity in this study are shown above.

## Results

### SH2scan Assay Principle and Compound Screening Pipeline

We developed SH2*scan* as a panel with 95 wildtype SH2 domain-containing constructs covering a total of 97 SH2 domains (>80% of the target class), as well as 7 mutant STAT SH2 domain-containing constructs (**Extended Data Table 1**). The platform uses a competition binding assay principle (**Fig. 2a and 2b**). Multiple binding reactions can be performed in parallel with a dose-response of a given competitor; these data are plotted, and a KD value is calculated from the curve fitting for the given protein-ligand interaction.

**Fig. 2.**
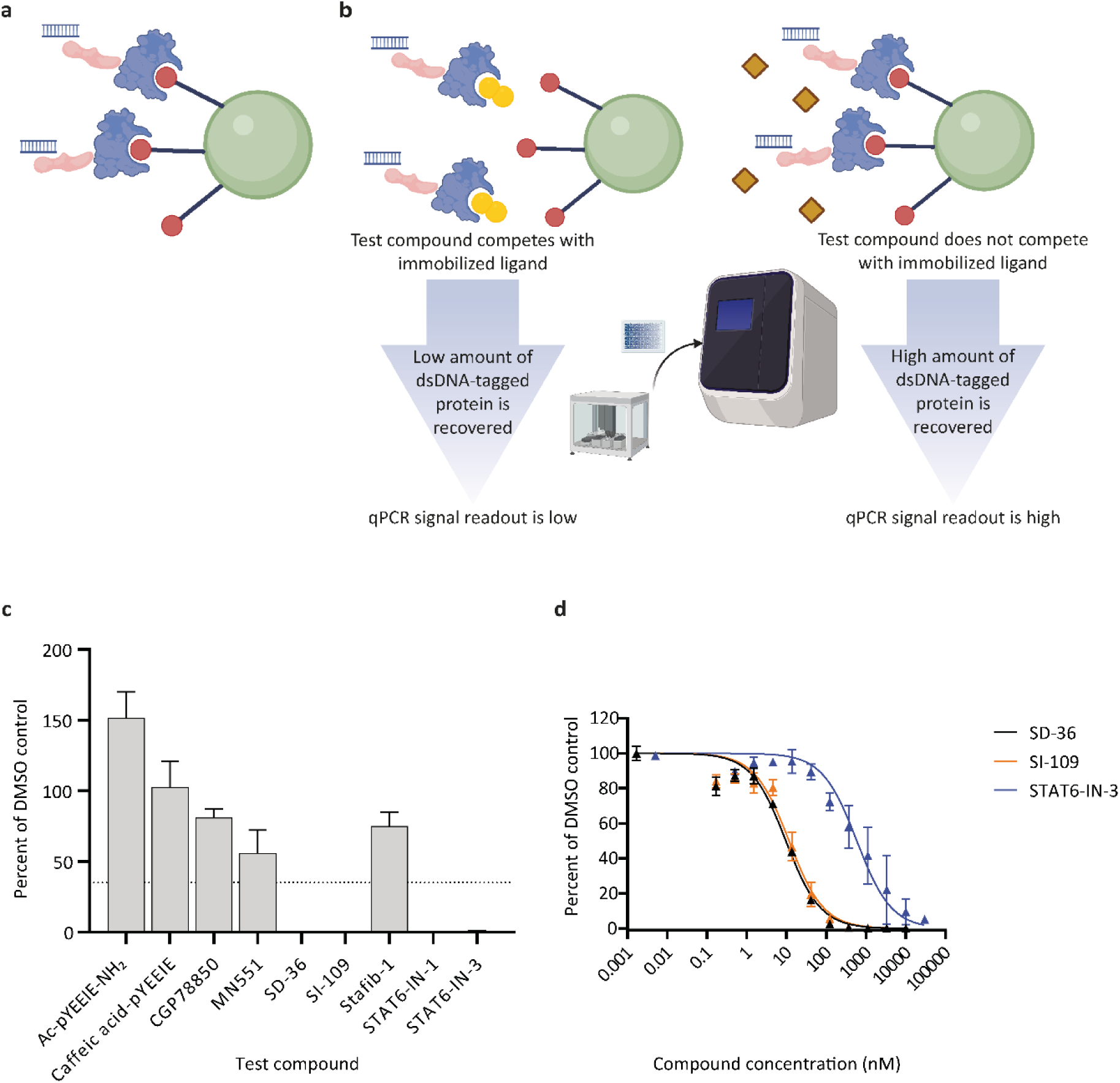
SH2scan assay principle for **measuring compound dissociation constants. a**, An SH2 domain- containing protein construct (blue) fused to the NFκB DNA binding domain (pink) is tagged with an exogenous double-stranded DNA (dsDNA) probe. This construct is incubated with a capture ligand (red) immobilized on magnetic beads (green). **b**, In the presence of a competitor compound (yellow), less tagged protein is captured on the beads. The protein remaining on the beads is eluted using a high concentration of sodium phenyl phosphate, a generic competitor of phosphopeptide binding to SH2 domains^9^. After elution, a lower qPCR signal is obtained at the end of the assay. In the presence of a non-competitor compound ligand (brown), more tagged protein is captured on the beads and a high qPCR signal is observed. **c**, Representative primary screening data for the STAT3 construct are shown. Compounds were tested at 10 μM and the hit cutoff for the compound screen was set at ≤35% of the average signal of the DMSO control wells for each construct (dashed line). Percent assay signal for each compound is expressed as the mean of at least two independent technical replicates from at least one independent experiment, ± standard deviation (exact numbers of replicates for each compound tested are shown in **Extended Data Table 3**). **d**, KD data for selected validated hits for the STAT3 construct are shown. Compounds were tested in dose-response, and the data points were fit to the Hill equation (see Methods). Data are presented as mean percent assay signal from four independent technical replicates collected over two independent experiments, ± standard error of the mean.

To examine the selectivity profiles of synthetic ligands across the SH2 domain target class, we chose 9 compounds from the literature to profile in our panel (**Fig. 1**). These compounds were expected to have a range of primary targets based on literature sources and included inhibitors/synthetic ligands (Ac-pYEEIE-NH_2_ ^9^ and caffeic acid-pYEEIE^10^ , targeting SRC family SH2 domains; CGP78850^11^, targeting the GRB2 SH2 domain; MN551^12^, targeting the SOCS2 SH2 domain; SI-109^13^, targeting the STAT3 SH2 domain; Stafib-1^14^, targeting the STAT5B SH2 domain; STAT6-IN-1 and STAT6-IN-3^15^, targeting the STAT6 SH2 domain) and a protein degrader (SD-36^13^, targeting the STAT3 SH2 domain). We profiled each of these compounds in a primary screen and then validated all hits in dose-response to extract KD values for each protein-ligand interaction. Representative data for the primary screen are shown in **Fig. 2c**, while sample dose-response data are shown in **Fig. 2d**.

### Primary Screening Data Visualization of Compound Binding Selectivity and Dose Response Validation of Primary Screening Hits

To visually assess the range of binding events detected for each compound tested in SH2*scan*, a phylogenetic tree was generated containing the 120 *Homo sapiens* SH2 domains, and for each compound, branch tips of the tree were assigned dots proportionally sized to the normalized percent of DMSO signal from the primary screening assay for each SH2 domain represented in the panel (**Fig. 3**).

**Fig. 3.**
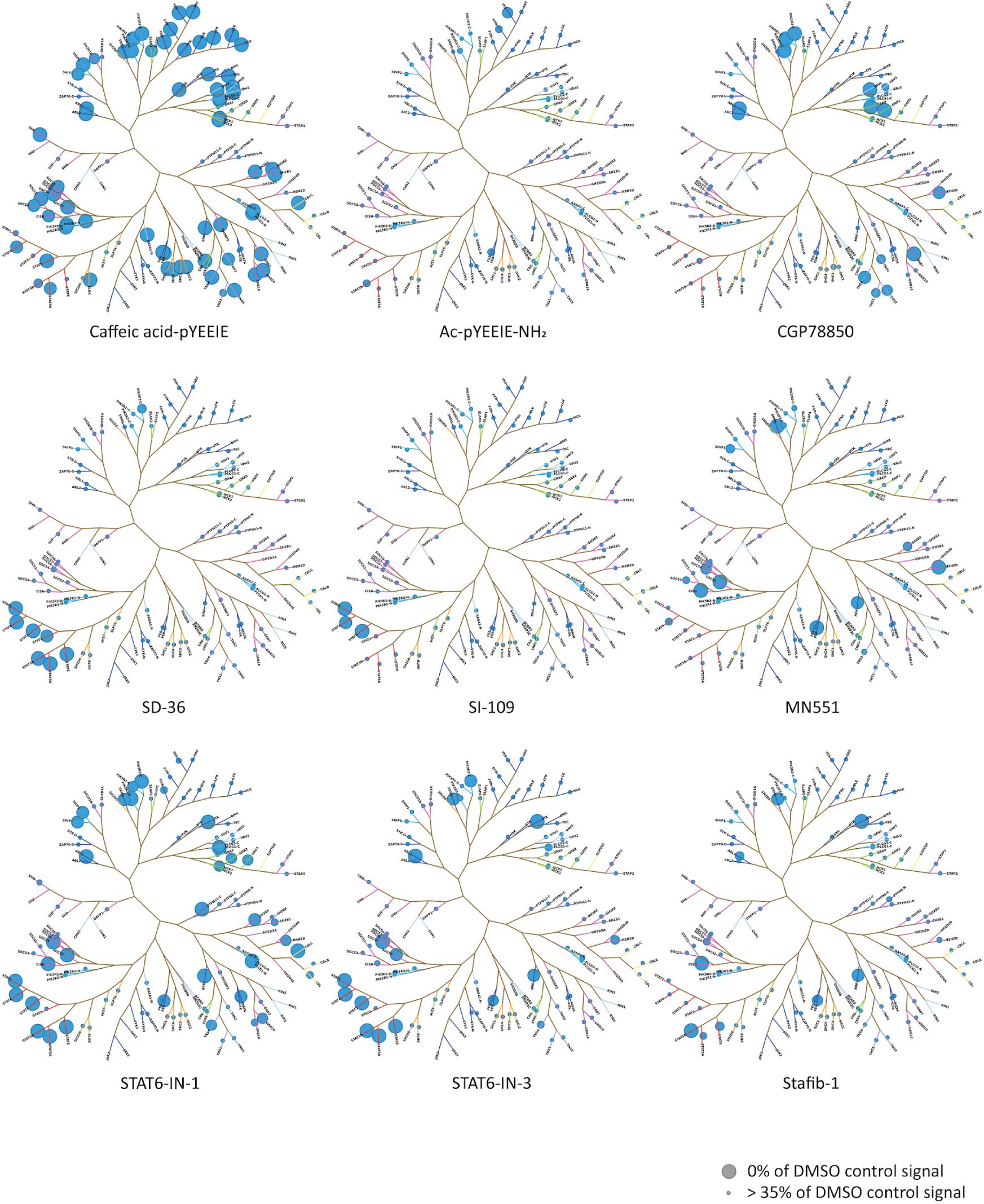
Primary screening across SH2scan reveals unique binding selectivity signatures for synthetic ligands. Percent of DMSO control values were measured for each compound tested in this study and mapped as dots on a phylogenetic tree containing all 120 canonical human SH2 domains. Each dot plotted in a diagram represents a mean value collected from at least two independent technical replicates collected over at least one independent experiment, ± standard deviation. Exact numbers of replicates for experiments are shown in **Extended Data Table 3**. The screening results for the two SH2 domains of SYK and ZAP70 are mapped as equivalently sized dots, given that the SH2 domains for these targets are expressed in tandem constructs, and any competition observed cannot be unequivocally assigned to one or the other SH2 domain.

Hits from the complete primary screen (**Extended Data Tables 2 and 3**) were validated in dose-response to obtain KD values (**Extended Data Tables 4 and 5**). Available literature values for these interactions are presented in **Extended Data Table 6**. Certain compound-SH2 domain target pairs which were previously described as having measured affinities in the literature^13^ that did not come up as hits in the primary screen were tested in dose-response, and in those cases where we measured KD values, these are indicated in **Extended Data Table 4**. Compounds which were above the hit cutoff in the primary screen that were validated as hits when tested in dose-response, and primary screening hits that did not confirm in the dose-response experiments, are also noted in this table.

Based on the KD values obtained for the validated SH2 domain-synthetic ligand binding hits, we were able to confirm engagement with the reported primary literature target(s) for each ligand, while also observing several off-target hits with varying levels of affinity for each compound (**Extended Data Tables 4 and 5**). One of the most striking examples of binding promiscuity we detected with SH2*scan* was for the semisynthetic phosphopeptide caffeic acid-pYEEIE, which was designed to bind to SRC family SH2 domains^10^. We confirmed with dose response data that this compound binds to 71 of the 102 constructs in the panel (∼70% of the constructs), with measured KD values as low as 0.53 nM (against the SRC SH2 domain family member FYN) and with 47 of the measured KDs in the submicromolar range. In contrast, the related phosphopeptide derivative Ac-pYEEIE-NH2 displayed a much more restricted binding profile, with measured affinities detected only for the constructs FES (KD = 4200 nM), FYN (KD = 3300 nM), PIK3R3(SH2dom.C-term. (denotes SH2 domain closest to C-terminus of protein)) (KD = 14900 nM) and YES (KD = 3800 nM).

For the GRB2 inhibitor CGP78850^11^, which targets the SH2 domain of this protein, we detected a strong on-target KD of 3.4 nM, as well as for the related SH2 domain family member GRAP (KD = 3.3 nM) and for the more distally related GRB7 (KD = 5.2 nM). Off-target binding events of note also included PIK3R1(SH2dom.C-term.) and PIK3R3(SH2dom.C-term.) (KD = 150 nM and 48 nM, respectively), TNS1, TNS2, TNS3 and TNS4 (KD = 408 nM, 1200 nM, 620 nM and 508 nM, respectively) and VAV1 and VAV3 (KD = 260 nM and 860 nM, respectively).

The STAT3 binding ligand SI-109 and SD-36^13^, a cereblon (CRBN)-recruiting PROTAC incorporating the SH2 domain binding moiety of SI-109, showed best binding to STAT3, with the KD for SI-109 being 15 nM and for SD-36 being 21 nM. The selectivity profile for these compounds was relatively narrow compared to other SH2-domain targeting compounds, with lower binding affinities detected for STAT1, STAT2, STAT4, STAT5A, STAT5B and STAT6 for both compounds, and a weak binding affinity detected for SOCS4 for SD-36 (KD = 5900 nM).

The SH2 domain-targeting covalent ligand of SOCS2, MN551^12^, had an on-target KD of 400 nM. We observed relatively strong interactions of the compound with SOCS4 (KD = 64 nM) and CISH (KD = 46 nM). Weak KDs in the micromolar range were observed for the compound for the targets HSH2D, LNK (SH2B3), PIK3R3 (SH2dom.C-term.) and SOCS7.

STAT6-IN-1 and STAT6-IN-3^15^, which bind to the SH2 domain of STAT6, showed strong on-target KD values of 0.68 nM and 8.6 nM, respectively, and relatively strong binding affinities for the other STAT SH2 domains. Interestingly, both compounds showed a strong off-target binding affinity for the SOCS4 construct (KDs of 13 and 10 nM, respectively). While a similar constellation of micromolar-range off-target binding affinities were observed for both compounds, additional weak binding events were observed for STAT6-IN-1 that were not present for STAT6-IN-3 for the targets CBL, CRK, CSK, GADS, GRB14, GRB2, ITK, LNK (SH2B3), MATK, PLCG2 (SH2dom.C-term.), PLCG2 (SH2dom.N-term. (denotes SH2 domain closest to N-terminus of protein)), PTPN11 (SH2dom.C-term.), SHC3, SHIP1, SHIP2, TNS4 and YES.

The STAT5B-selective inhibitor Stafib-1^14^, showed highest binding affinity to its reported target in our assay system (KD = 1800 nM), and micromolar range off-target affinities against the targets CISH, FER, PTPN6 (SH2dom.N-term.), SOCS4, STAT1, STAT2, STAT5A and STAT6.

## Discussion

We observed a wide range of affinities (picomolar to micromolar KD values) and selectivity profiles for the compounds tested in SH2*scan*. While some compounds tested showed relatively narrow selectivity profiles (such as SD-36, SI-109 and MN551), others (such as caffeic acid-pYEEIE, STAT6-IN-1 and STAT6-IN-3) showed promiscuity in terms of their SH2 domain binding. All 9 compounds tested in the platform showed varying degrees of off-target binding to SH2 domains other than their primary targets. These findings demonstrate the utility of compound screening across SH2 domains to identify off-target binding events early in the drug discovery process, to reduce the possibility of undesirable pharmacological effects.

We observed the best binding affinity in our assay system for the reported primary target for each of the 9 compounds tested. In addition, we revealed additional off-target binding events that were previously reported (such as the binding of MN551 to CISH^12^ and SI-109/SD-36 binding to STAT family SH2 domains^13^), as well as new off-target events (such as the strong interaction of STAT6-IN-1 and STAT6-IN-3 with SOCS4). Given that SOCS protein-containing E3 ligase complexes are thought to be involved in the suppression of JAK-STAT pathway effectors via the ubiquitin-proteosome system^16^, the binding of these two STAT inhibitors to the SH2 domain of SOCS4 might interfere with their intended anti-inflammatory effects. These interesting results merit follow-up in future studies. With respect to the structure of these two inhibitors, STAT6-IN-1 differs from STAT6-IN-3 by the presence of a single additional methyl group. From our data, this seems to afford a greater on-target potency for STAT6-IN-1 as compared to STAT6-IN-3, at the expense of a loss of selectivity (there were 17 additional targets which came up as weak binders to STAT6-IN-1 during the dose response validation process). This highlights the importance of considering off-target effects during the structure-activity relationship (SAR) process when designing selective compounds for SH2 domains: our panel provides the opportunity to track the selectivity of compounds in parallel with on-target potency optimization during compound series derivatization. This study has revealed that even small changes to the chemical structure of SH2 domain-targeting compounds can have profound effects on binding affinity and selectivity within the target class.

Many SH2 domain-containing proteins present in SH2*scan* have limited or non-existent biochemical and functional characterization in the scientific literature. This platform presents the opportunity for the screening of chemical matter for the development of new molecular probe tool compounds to interrogate the function of these SH2 domain-containing proteins. Furthermore, this platform can be used to dissect PPI networks involving SH2 domains at scale via profiling of phosphopeptides derived from proteins of interest. These applications can be used to discover new biological insight and actionable targets for therapeutic intervention.

In sum, this work presents SH2*scan* as an assay platform that can be leveraged at scale to both develop new therapeutics targeting SH2 domains as well as to uncover new PPI networks.

## Online Methods

### Small molecules and nucleic acids

Caffeic acid-pYEEIE, CGP78850, MN551, SD-36, SI-109, Stafib-1, STAT6-IN-1, and STAT6-IN-3 were purchased from MedChemExpress. The Ac-pYEEIE-NH2 phosphopeptide was custom synthesized and purchased from Biopeptide Co., Inc. The DNA probe containing the qPCR amplicon for tagging the NFkB fusion domain was custom synthesized and purchased from Thermo Fisher Scientific.

### Protein constructs and protein expression

Wild-type and mutant SH2-domain containing protein constructs were engineered as N-terminal fusions with the DNA-binding domain of NFkB (amino acids 35–36 fused to amino acids 41–359, with UniProt entry P19838 being used as a reference)^17^. Expression of SH2 domain-containing proteins was achieved through transient transfection of HEK293 cells. The resulting protein extracts were prepared using M-PER extraction buffer (Pierce Biotechnology), with added 150 mM NaCl, 10 mM DTT, cOmplete™ Protease Inhibitor Cocktail (Roche Diagnostics GmbH) and Phosphatase Inhibitor Cocktail Set II (Merck KGaA) according to the manufacturer’s guidelines. Phosphatase inhibitors were omitted for the non-phosphorylated (np) STAT3 mutant extracts: np-STAT3(N647I), np-STAT3(S614R), np-STAT3(D661Y), np-STAT3(Y640F), np-STAT3(N567K), and np-STAT3(E616del). These extracts were incubated at 30 °C for 45 min during the protein harvest protocol to allow for dephosphorylation of STAT3 phosphotyrosines by endogenous phosphatases. All other extracts for the STAT3 mutant constructs were performed as per the manufacturer’s instructions. Construct details are provided in **Extended Data Table 1**.

### Competition binding assays

Competition binding assays were adapted from protocols used for kinases, bromo domains and KRAS^18–23^. For liganded beads, preparation was performed as follows: streptavidin-coated magnetic beads (Thermo Fisher Scientific) were incubated with biotinylated phosphopeptide capture ligands which were custom synthesized (Biopeptide Co., Inc) at 25 °C for 30 min. In order to remove the unbound affinity ligand and to reduce the nonspecific binding of proteins in the cell lysate, excess biotin (125 mM) was subsequently introduced to the ligand-coated beads. The beads were then washed with a blocking buffer containing SeaBlock (Pierce Biotechnology), 1% BSA and 0.05% Tween 20.

The binding reactions were performed with protein extracts containing the DNA-tagged SH2 domain-containing construct, affinity ligand-coated beads and a given small molecule test compound in a binding buffer (10 mM HEPES, 50 mM NaCl, 1 mM EDTA, 0.01% Tween 20) in a deep well, natural polypropylene 384-well plate, catalog number 784201 (Greiner Bio-One), in a final volume of 19.7 μL. No purification of SH2 domain-containing constructs was conducted prior to adding the protein extracts to the reaction mixture. The extracts were diluted 10,000-fold in the final reaction, resulting in a DNA-tagged construct concentration of less than 0.1 nM. Binding assay reactions were incubated at 25 °C with shaking for 1 h. After the incubation period, the affinity ligand-coated beads were thoroughly washed with the buffer used in the binding step to remove any non-specifically bound protein from the beads. The remaining bead-bound protein was then eluted from the beads via resuspension and incubation (30 min, 25°C) in an elution buffer of 20 mM sodium phenyl phosphate dibasic hydrate (Sigma-Aldrich) dissolved in the binding buffer. Quantitative PCR was then used to determine the concentration of SH2-domain containing protein constructs in the eluates. Primary screens were conducted using 10 μM of a given test compound. Hits were then validated in dose-response to extract KD values for each test compound and were determined using a series of 11-point, 3-fold serial dilutions.

The concentration of the capture ligand present on the magnetic beads was optimized in each assay to ensure accurate determination of true thermodynamic KD values for test compounds, as previously described in detail for this type of assay^21^. The complete primary screening data set is shown in **Extended Data Tables 2 and 3**, and the complete dose-response validation data set for this study is presented in **Extended Data Tables 4 and 5**.

### Curve fitting for competitive binding assays

Thermodynamic dissociation constants (KDs) were derived for each competitor-SH2 domain-containing construct interaction by fitting the data to a standard dose-response curve using the Hill equation:

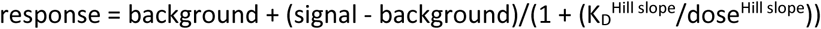

The Hill slope was set to -1 and a nonlinear least-square fit using the Levenberg-Marquardt algorithm was performed to generate the final curves.

### Generation of SH2 domain phylogenies

Phylogenetic trees were generated based on the sequence for each canonical *Homo sapiens* SH2 domain. Protein sequences for the SH2 domain regions were mined from NCBI reference sequences, aligned using MUSCLE^24^ in SeaView v.5.0.5^25^ and were further refined using GBlocks^26^ with parameters allowing for the least amount of stringency for retention^27^. Phylogenetic analysis was conducted on the Cyberinfrastructure for Phylogenetic Research (CIPRES) Science Gateway (www.phylo.org^28^) using the RAxML software v.8.2.12^29^ with the LG evolutionary and GTRGAMMA models^30^. Branch support was estimated by bootstraps with 700 replications and the constructed phylogram was edited using ITOL (itol.embl.de). SH2 protein nomenclature was determined using UNIPROT (uniprot.org). Accession numbers for individual proteins are available in **Extended Data Table 1**.

### Analysis of primary screening and dose-response data

Independent replicates for each primary screening and dose-response validation data set are noted in **Extended Data Table 3** and **Extended Data Table 5**, respectively. For each independent target assessed in both primary screening and dose-response formats, an assay signal was calculated from the vehicle control (DMSO) and background from a positive control compound. Passing assay metrics were defined as a signal to background ratio of ≥3 and a Z score of ≥0.3 calculated in Spotfire™ (Tibco). For primary screening experiments, the percent of control (POC) for each replicate per compound was calculated by dividing the assay signal of the test compound by that of the vehicle control. Positive hits were defined as ≤35% POC. For dose-response experiments, KD values were calculated for each individual replicate curve.

### Mapping of primary screening values onto phylogenetic trees

Phylogenetic trees were mapped in MS Paint and the resulting X,Y coordinates loaded into Spotfire™(Tibco) along with the tree image described above. Primary screening data was mapped onto the resulting tree template by subtracting the average POC for the test compound from 100 in order to generate dots directly proportional to the strength of each hit. These values were then loaded into Spotfire™(Tibco) along with the corresponding X,Y coordinates and used to generate a spot layer overlay onto the phylogenetic tree.

## Data availability

Chemical structures in Fig. 1 were generated using ChemDraw. Images in Fig.2a and Fig.2b were generated using Biorender.com and in Fig.2c and Fig.2d using GraphPad Prism. Construct details, primary screening data and dose-response data for this study are presented in Extended Data Tables 1-5. Any other data that support the conclusions of this paper are available upon request.

### Acknowledgements

We thank members of Eurofins DiscoverX, LLC for reviewing the manuscript and providing feedback.

## Author contributions

Design of the clones and capture ligand molecules was performed by L.M.G.L. and J.A.B. Construct cloning and protein lysate generation were performed by L.M.G.L., A.M.C., T.D.L., G.M.K., M.K., M.N., J.A.N. and J.A.B. Assay development and optimization was conducted by L.M.G.L., A.M.C., T.D.L., G.M.K., M.K., M.N., J.A.N. and J.A.B. Compound screening was performed by J.K.W.D. The phylogenetic tree used for compound selectivity graphics was generated by A.M.C. Mapping of screening data to the phylogenetic tree diagram was conducted by J.K.W.D. and G.P. Data analysis for compound screening data was performed by J.K.W.D., G.P., N.B.S. and J.A.B. Experimental conceptualization and project supervision were conducted by J.A.B. The paper was written by L.M.G.L., J.K.W.D., A.M.C., T.D.L., G.P. and J.A.B. All authors reviewed the manuscript.

---

**Extended Data Table 1.** SH2*scan* protein construct details. Shown are the gene names, construct names, NCBI protein accession numbers, amino acid ranges and mutation status for all proteins in SH2*scan*. SH2dom.C-term. indicates a construct containing the SH2 domain closer to the C-terminus of the full-length protein when more than one SH2 domain is present in the protein. SH2dom.N-term. indicates a construct containing the SH2 domain closer to the N-terminus of the full-length protein when more than one SH2 domain is present in the protein. SH2dom1.dom2. signifies a construct containing two tandem SH2 domains (these engage a doubly phosphorylated phosphopeptide capture ligand). Non-phosphorylated constructs are harvested from protein lysates that are not treated with phosphatase inhibitors, as described in the Methods section.

| Gene Name | Construct Name | NCBI protein accession number | Amino acid range | Mutant |
| --- | --- | --- | --- | --- |
| ABL1 | ABL1 | NP_009297.2 | S140/Y251 | No |
| ABL2 | ABL2 | NP_009298.1 | V165/G273 | No |
| APS (SH2B2) | APS (SH2B2) | NP_001346157.1 | E409/Q518 | No |
| BLK | BLK | NP_001706.2 | V113/A224 | No |
| BLNK | BLNK | NP_037446.1 | G317/S456 | No |
| BRK | BRK | NP_005966.1 | S75/E174 | No |
| CBL | CBL | NP_005179.2 | P47/G351 | No |
| CBLB | CBLB | NP_001308715.1 | A67/F454 | No |
| CBLC | CBLC | NP_036248.3 | M1/G323 | No |
| CISH | CISH | NP_037456.5 | K76/L275 | No |
| CRK | CRK | NP_058431.2 | D6/G124 | No |
| CRKL | CRKL | NP_005198.1 | M1/Y105 | No |
| CSK | CSK | NP_004374.1 | M80/M173 | No |
| DAPP1 | DAPP1 | NP_055210.2 | Q31/R131 | No |
| FER | FER | NP_005237.2 | K453/K552 | No |
| FES | FES | NP_001996.1 | V451/K551 | No |
| FGR | FGR | NP_005239.1 | S138/A255 | No |
| FRK | FRK | NP_002022.1 | A106/V213 | No |
| FYN | FYN | NP_002028.1 | W149/K248 | No |
| GADS | GADS | NP_004801.1 | W58/H155 | No |
| GRAP | GRAP | NP_006604.1 | P59/E151 | No |
| GRB10 | GRB10 | NP_001357938.1 | I536/R640 | No |
| GRB14 | GRB14 | NP_004481.2 | I433/R537 | No |
| GRB2 | GRB2 | NP_002077.1 | P59/I151 | No |
| GRB7 | GRB7 | NP_001025173.1 | P415/L532 | No |
| HCK | HCK | NP_002101.2 | L139/K245 | No |
| HSH2D | HSH2D | NP_116244.1 | G24/D129 | No |
| ITK | ITK | NP_005537.3 | N232/C339 | No |
| LCK | LCK | NP_005347.3 | A119/T226 | No |
| LNK (SH2B3) | LNK (SH2B3) | NP_005466.1 | D356/V465 | No |
| LYN | LYN | NP_002341.1 | S115/P229 | No |
| MATK | MATK | NP_002369.2 | L118/G216 | No |
| MIST | MIST | NP_443196.2 | W291/T422 | No |
| NCK1 | NCK1 | NP_001278928.1 | P281/S377 | No |
| NCK2 | NCK2 | NP_003572.2 | E284/Q380 | No |
| PIK3R1 | PIK3R1(SH2dom.C-term.) | NP_852664.1 | L616/R724 | No |
| PIK3R1 | PIK3R1(SH2dom.N-term.) | NP_852665.1 | G21/Q135 | No |
| PIK3R2 | PIK3R2(SH2dom.C-term.) | NP_005018.2 | D610/R717 | No |
| PIK3R2 | PIK3R2(SH2dom.N-term.) | NP_005018.2 | A317/Q432 | No |
| PIK3R3 | PIK3R3(SH2dom.C-term.) | NP_001290357.1 | N366/C477 | No |
| PIK3R3 | PIK3R3(SH2dom.N-term.) | NP_001290357.1 | S74/E188 | No |
| PLCG1 | PLCG1(SH2dom.C-term.) | NP_002651.2 | H663/E759 | No |
| PLCG1 | PLCG1(SH2dom.N-term.) | NP_002651.2 | E533/A662 | No |
| PLCG2 | PLCG2(SH2dom.C-term.) | NP_002652.2 | H641/E738 | No |
| PLCG2 | PLCG2(SH2dom.N-term.) | NP_002652.2 | E516/P640 | No |
| PTPN11 | PTPN11(SH2dom.C-term.) | NP_001317366.1 | T108/R220 | No |
| PTPN11 | PTPN11(SH2dom.N-term.) | NP_001317366.1 | M1/P107 | No |
| PTPN6 | PTPN6(SH2dom.C-term.) | NP_536859.1 | T106/R217 | No |
| PTPN6 | PTPN6(SH2dom.N-term.) | NP_536859.1 | M1/D104 | No |
| RASA1 | RASA1(SH2dom.C-term.) | NP_002881.1 | G340/Q444 | No |
| RASA1 | RASA1(SH2dom.N-term.) | NP_002881.1 | T174/D280 | No |
| SH2B (SH2B1) | SH2B (SH2B1) | NP_001139267.1 | D519/S628 | No |
| SH2D1A | SH2D1A | NP_002342.1 | M1/K104 | No |
| SH2D1B | SH2D1B | NP_444512.2 | M1/R103 | No |
| SH2D6 | SH2D6 | NP_001381392.1 | S213/P335 | No |
| SH3BP2 | SH3BP2 | NP_001139328.1 | E503/R618 | No |
| SHB | SHB | NP_003019.2 | E397/L509 | No |
| SHC1 | SHC1 | NP_001123512.1 | A481/L584 | No |
| SHC2 | SHC2 | NP_036567.2 | E479/P582 | No |
| SHC3 | SHC3 | NP_058544.3 | E492/Q594 | No |
| SHC4 | SHC4 | NP_976224.3 | K518/K630 | No |
| SHF | SHF | NP_001288097.2 | L336/L441 | No |
| SHIP1 | SHIP1 | NP_001017915.1 | M1/P112 | No |
| SHIP2 | SHIP2 | NP_001558.3 | S20/V117 | No |
| SLAP2 | SLAP2 | NP_115590.1 | A88/L193 | Np |
| SLP76 | SLP76 | NP_005556.1 | N404/T527 | No |
| SOCS1 | SOCS1 | NP_003736.1 | S60/I211 | No |
| SOCS2 | SOCS2 | NP_001257400.1 | Q32/V198 | No |
| SOCS3 | SOCS3 | NP_003946.3 | M1/L225 | No |
| SOCS4 | SOCS4 | NP_955453.1 | L274/E437 | No |
| SOCS6 | SOCS6 | NP_004223.2 | V361/S499 | No |
| SOCS7 | SOCS7 | NP_055413.2 | S451/F576 | No |
| SRC | SRC | NP_938033.1 | Q147/T250 | No |
| SRMS | SRMS | NP_543013.1 | D117/M213 | No |
| STAP1 | STAP1 | NP_036240.1 | Y168/H293 | No |
| STAP2 | STAP2 | NP_060190.2 | V138/D247 | No |
| STAT1 | STAT1 | NP_009330.1 | L136/S710 | No |
| STAT2 | STAT2 | NP_005410.1 | G130/L706 | No |
| STAT3 | STAT3 | NP_001356441.1 | G127/I722 | No |
| STAT3 | STAT3(D661Y) -nonphosphorylated | NP_001356441.1 | G127/I722 | D661Y |
| STAT3 | STAT3(E616del)-nonphosphorylated | NP_001356441.1 | G127/I722 | E616del |
| STAT3 | STAT3(N567K)-nonphosphorylated | NP_001356441.1 | G127/I722 | N567K |
| STAT3 | STAT3(N647I)-nonphosphorylated | NP_001356441.1 | G127/I722 | N647I |
| STAT3 | STAT3(S614R)-nonphosphorylated | NP_001356441.1 | G127/I722 | S614R |
| STAT3 | STAT3(Y640F) -nonphosphorylated | NP_001356441.1 | G127/I722 | Y640F |
| STAT4 | STAT4 | NP_001230764.1 | S137/S702 | No |
| STAT5A | STAT5A | NP_001275647.1 | S128/D712 | No |
| STAT5B | STAT5B | NP_036580.2 | S128/D717 | No |
| STAT5B | STAT5B(N642H) | NP_036580.2 | S128/D717 | N642H |
| STAT6 | STAT6 | NP_003144.3 | M119/Q662 | No |
| SYK | SYK(SH2dom1.dom2.) | NP_003168.2 | S9/I262 | No |
| TEC | TEC | NP_003206.2 | T235/T355 | No |
| TNS1 | TNS1 | NP_072174.3 | D1458/D1574 | No |
| TNS2 | TNS2 | NP_056134.2 | D1145/D1259 | No |
| TNS3 | TNS3 | NP_073585.8 | D1167/D1284 | No |
| TNS4 | TNS4 | NP_116254.4 | D444/E558 | No |
| TXK | TXK | NP_003319.2 | N140/S251 | No |
| VAV1 | VAV1 | NP_005419.2 | H660/T772 | No |
| VAV2 | VAV2 | NP_001127870.1 | S663/S773 | No |
| VAV3 | VAV3 | NP_006104.4 | V661/S772 | No |
| YES | YES | NP_005424.1 | P149/T262 | No |
| ZAP70 | ZAP70(SH2dom1.dom2.) | NP_001070.2 | D3/N256 | No |

**Extended Data Table 2.**
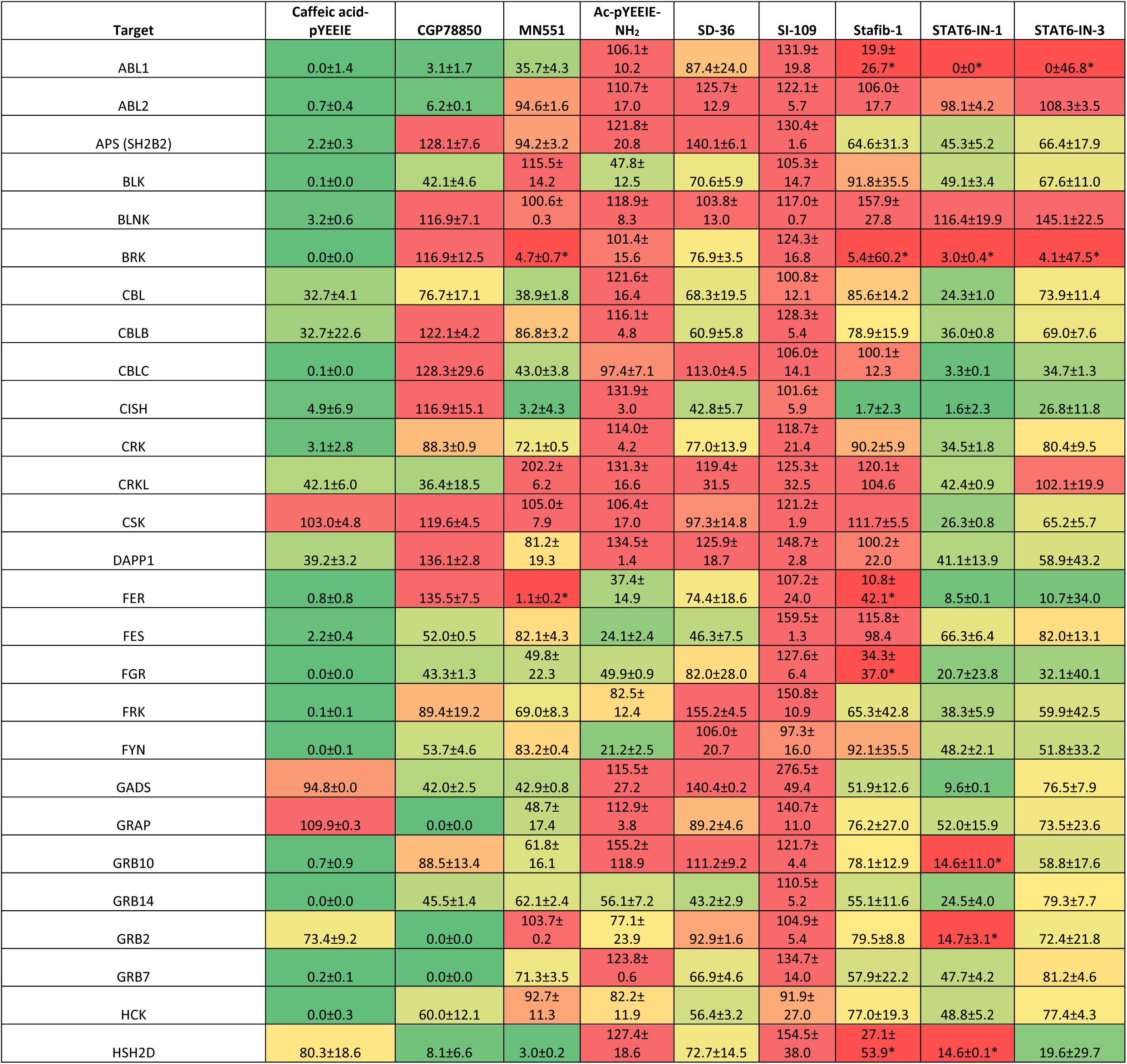
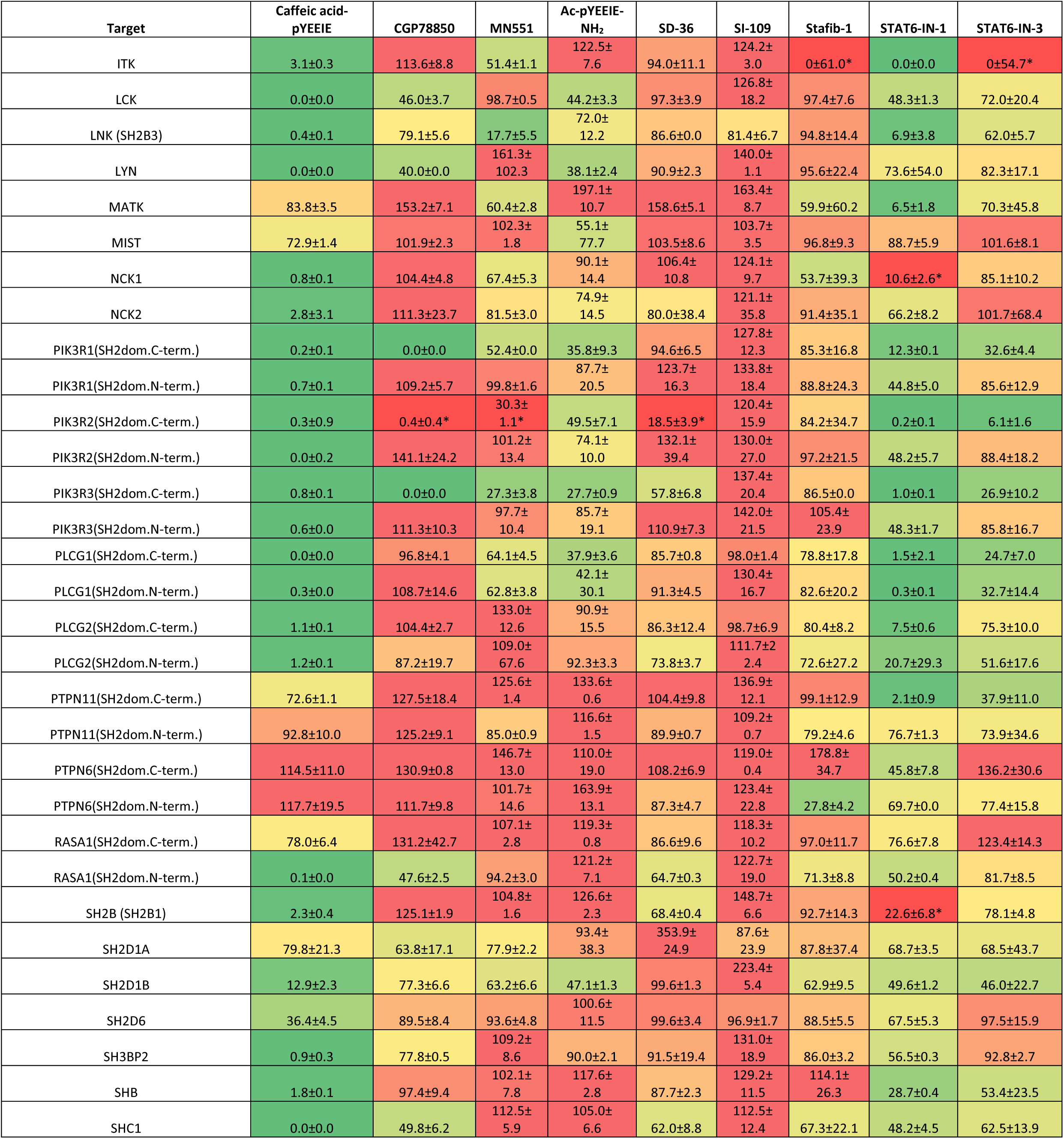
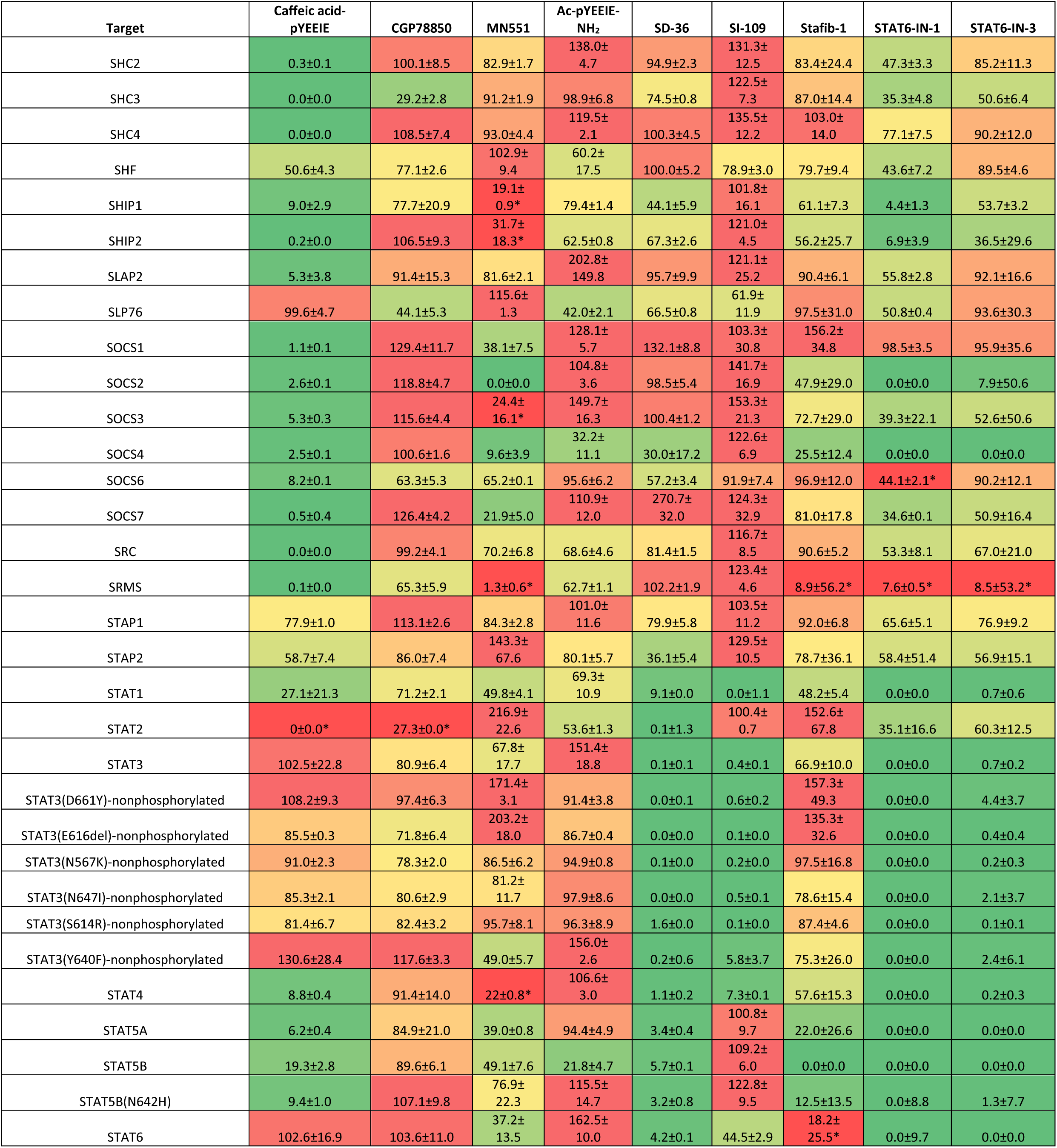
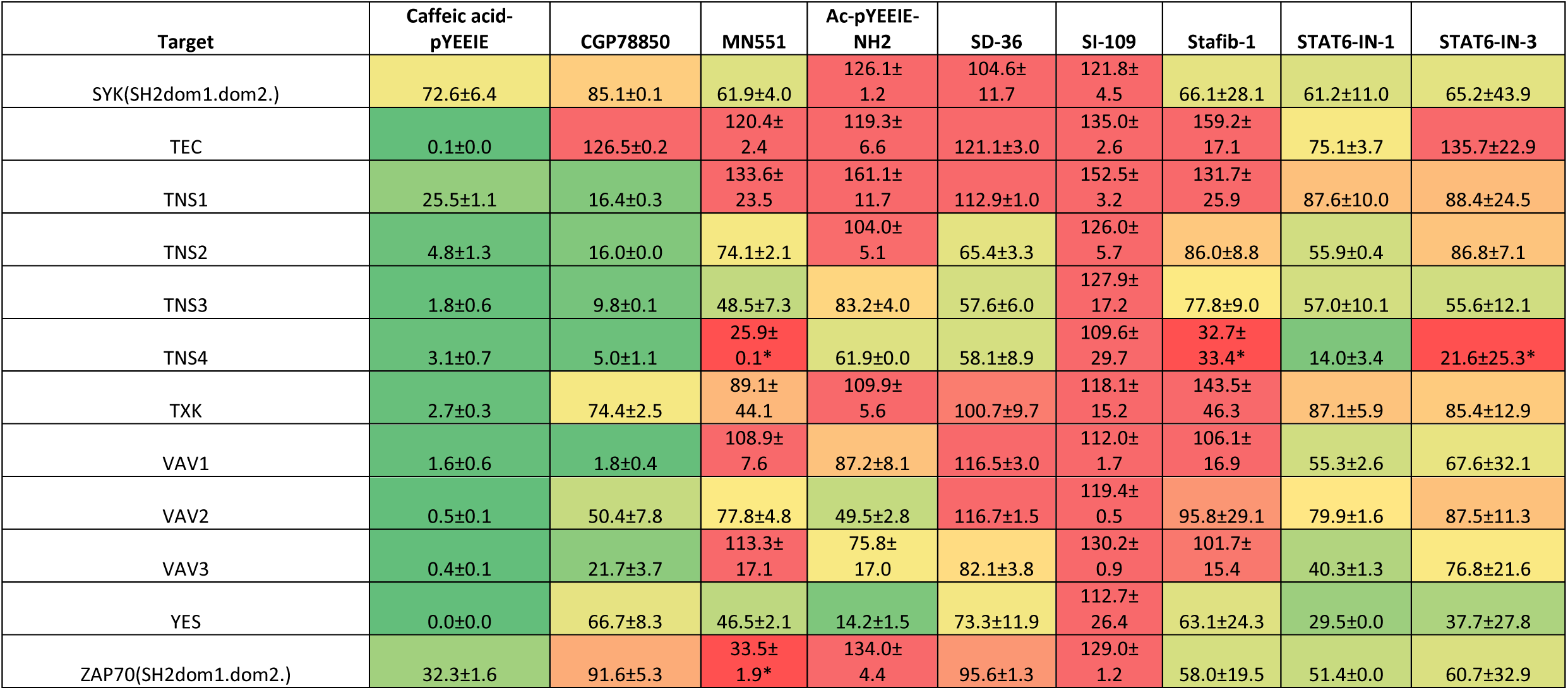
Primary screening data for the 9 compounds tested in SH2*scan*. Data is expressed as the mean of the normalized percent of the DMSO vehicle control, ± standard deviation. The number of independent replicates and independent experiments for the shown data are presented in **Extended Data Table 3**. False positives are marked with an asterisk.

**Extended Data Table 3.**
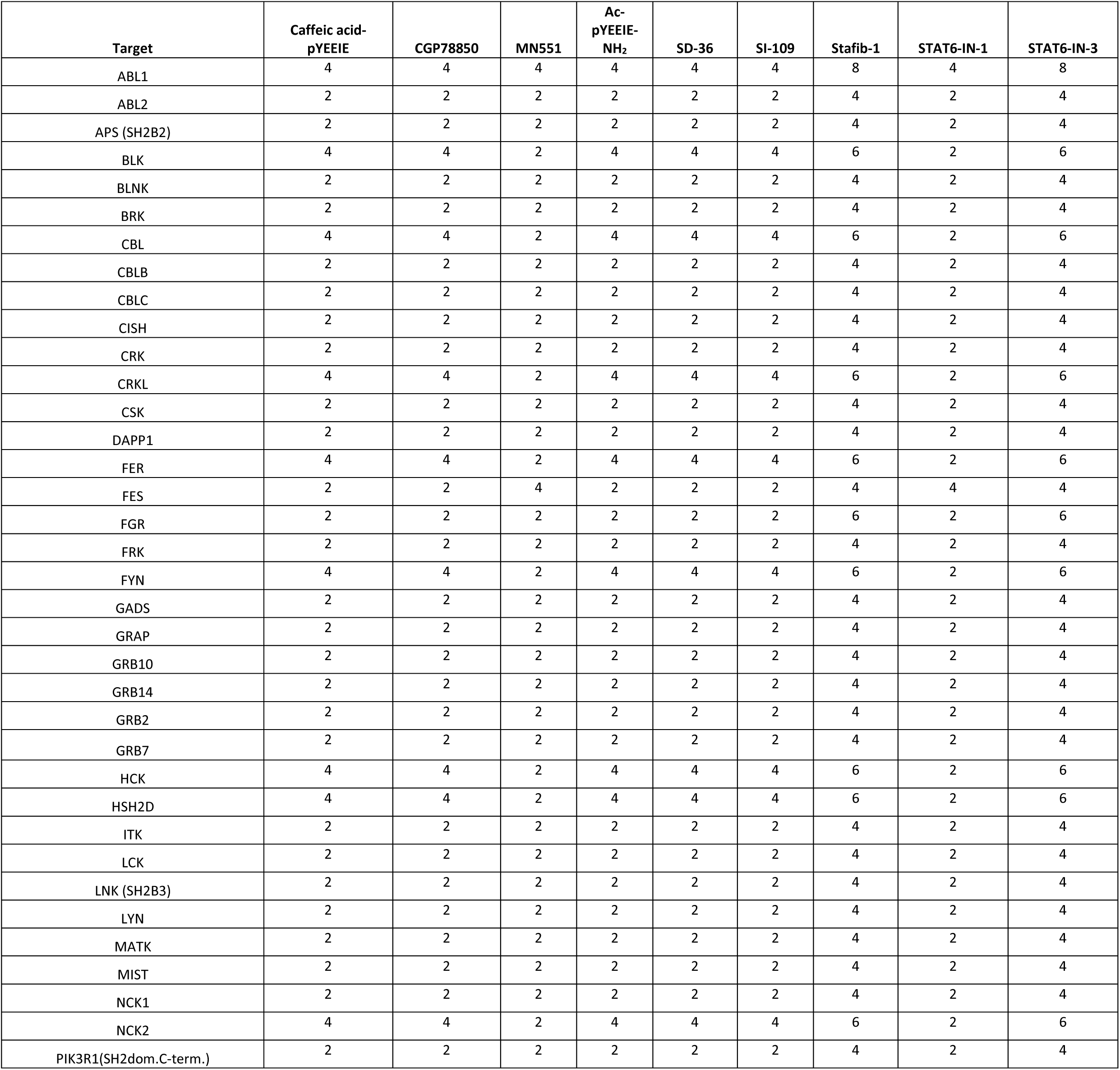
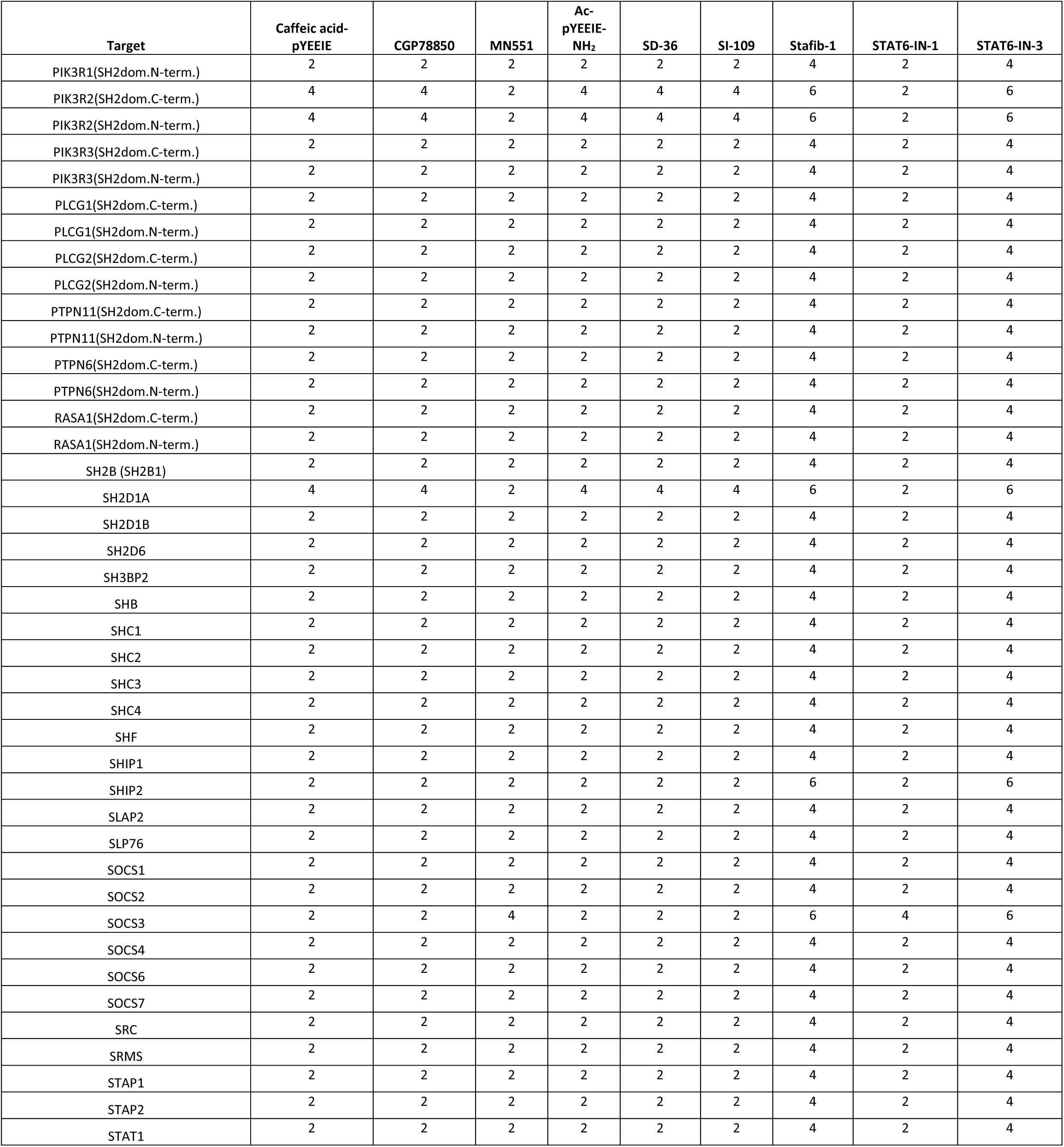
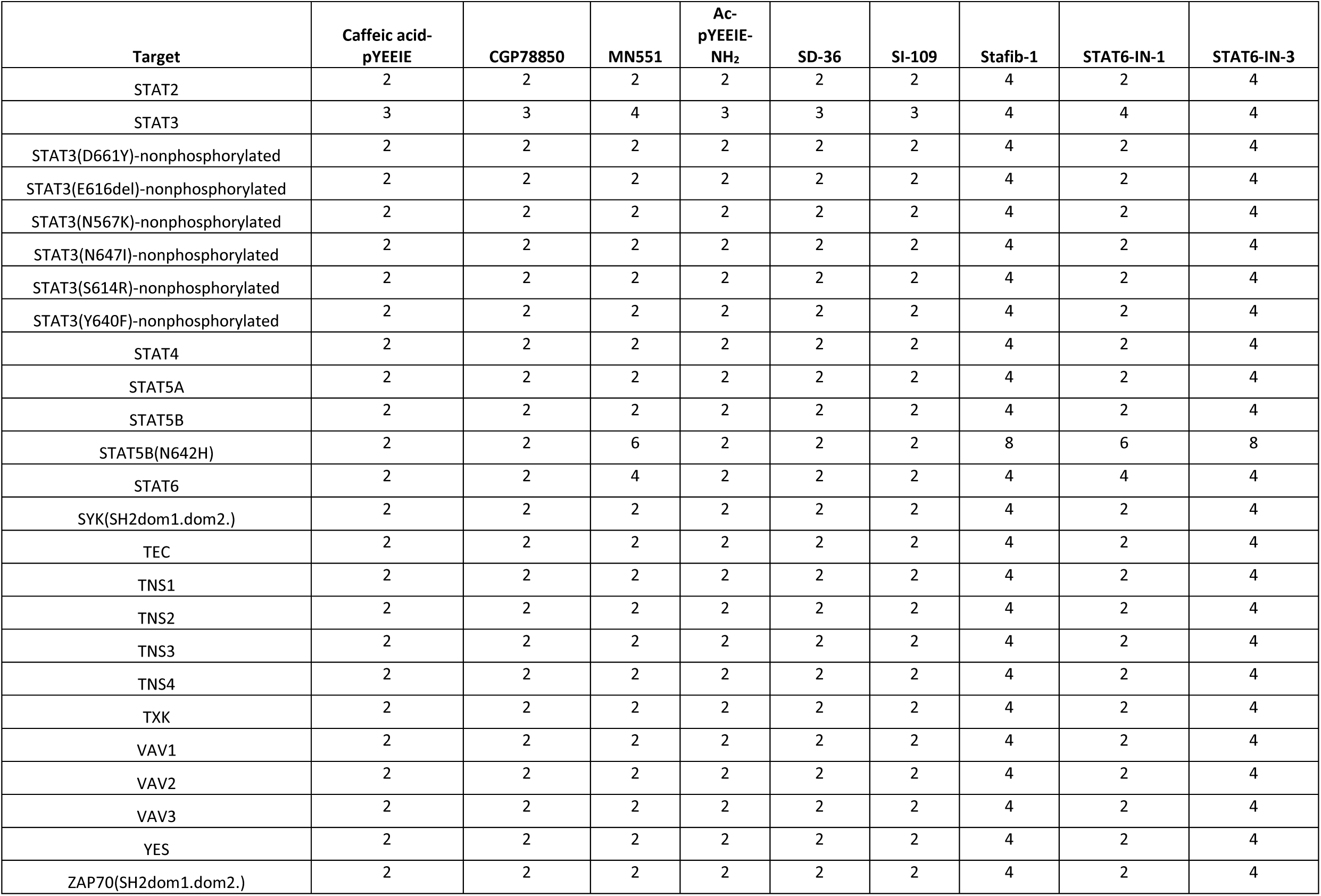
Replicate numbers for primary screening data for the 9 compounds tested in SH2*scan*. Independent replicates numbers (N) for each primary screening value presented in **Extended Data Table 2** are shown below and were generated from at least one independent experiment.

**Extended Data Table 4.**
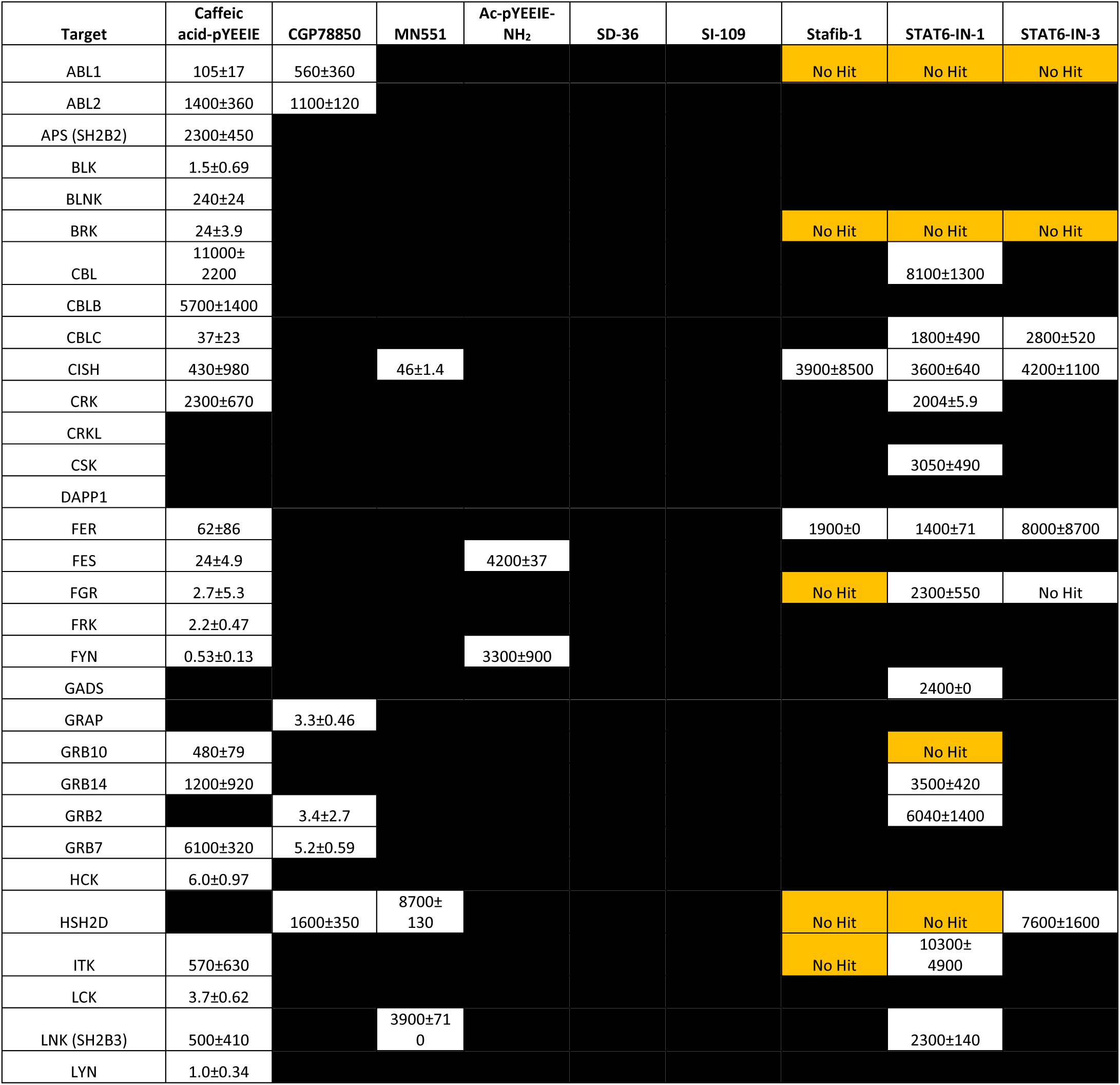
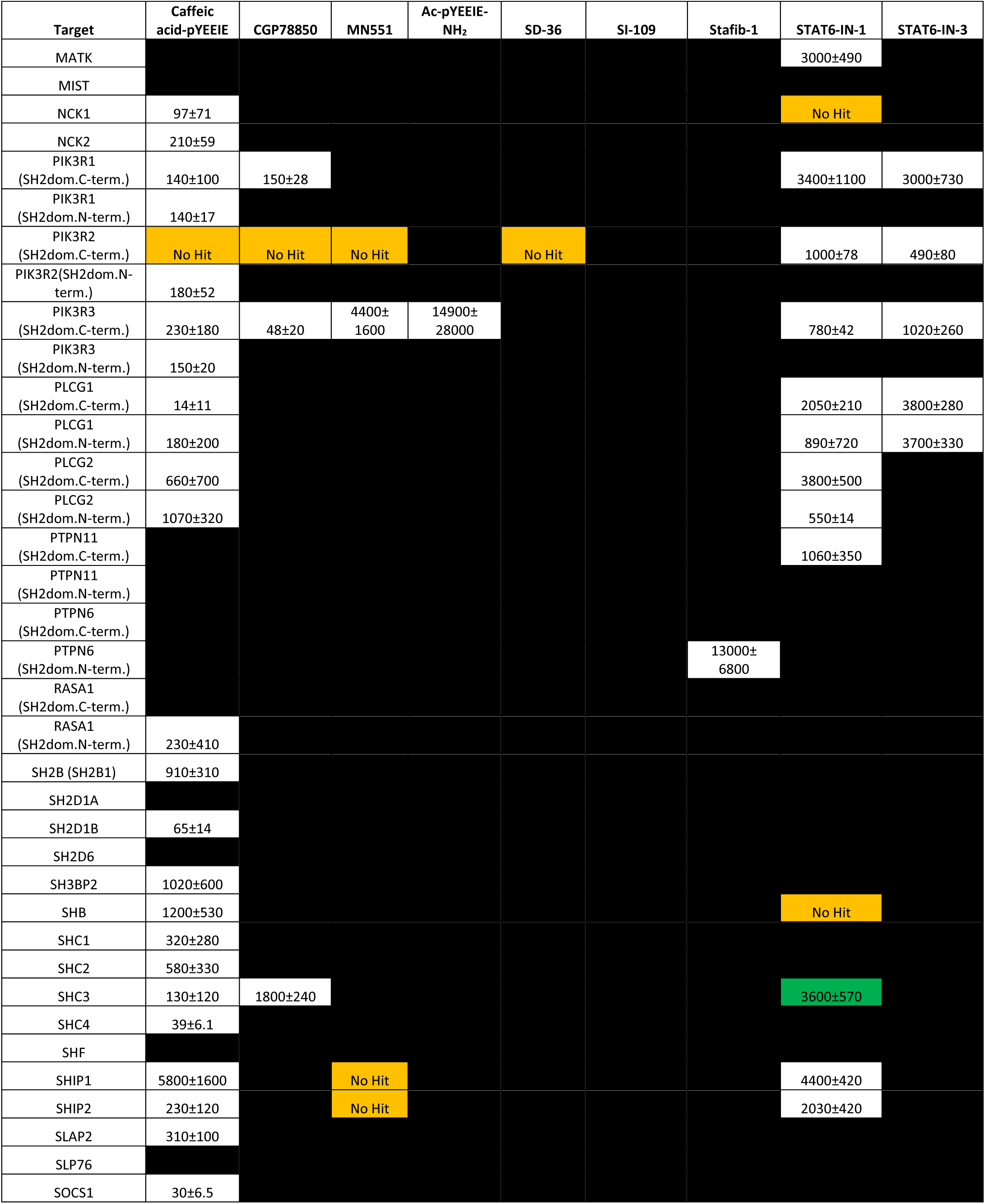
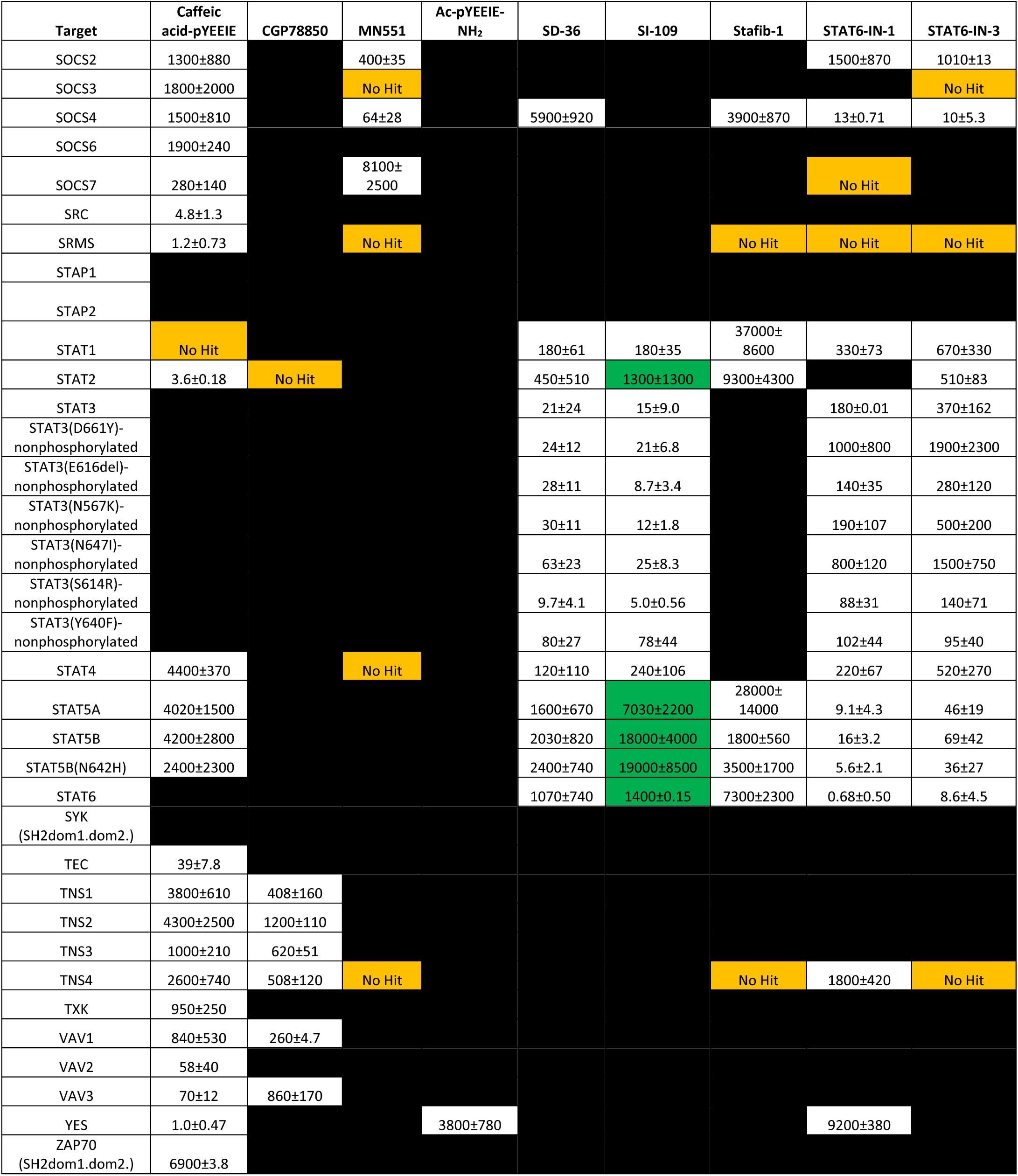
Dissociation constant values for the 9 compounds tested in SH2*scan*. Mean KD values collected in this study are listed below in nM, ± standard deviation. The number of independent replicates and independent experiments for the shown data are presented in **Extended Data Table 5**. False positives are colored in orange and are denoted with “no hit”. False negatives are colored in green. The false positive rate for the study was calculated as 3.8% (35 false positives/918 total tested conditions x 100). The false negative rate for the study was calculated as 0.9% (9 false negatives/918 total tested conditions x 100).

**Extended Data Table 5.**
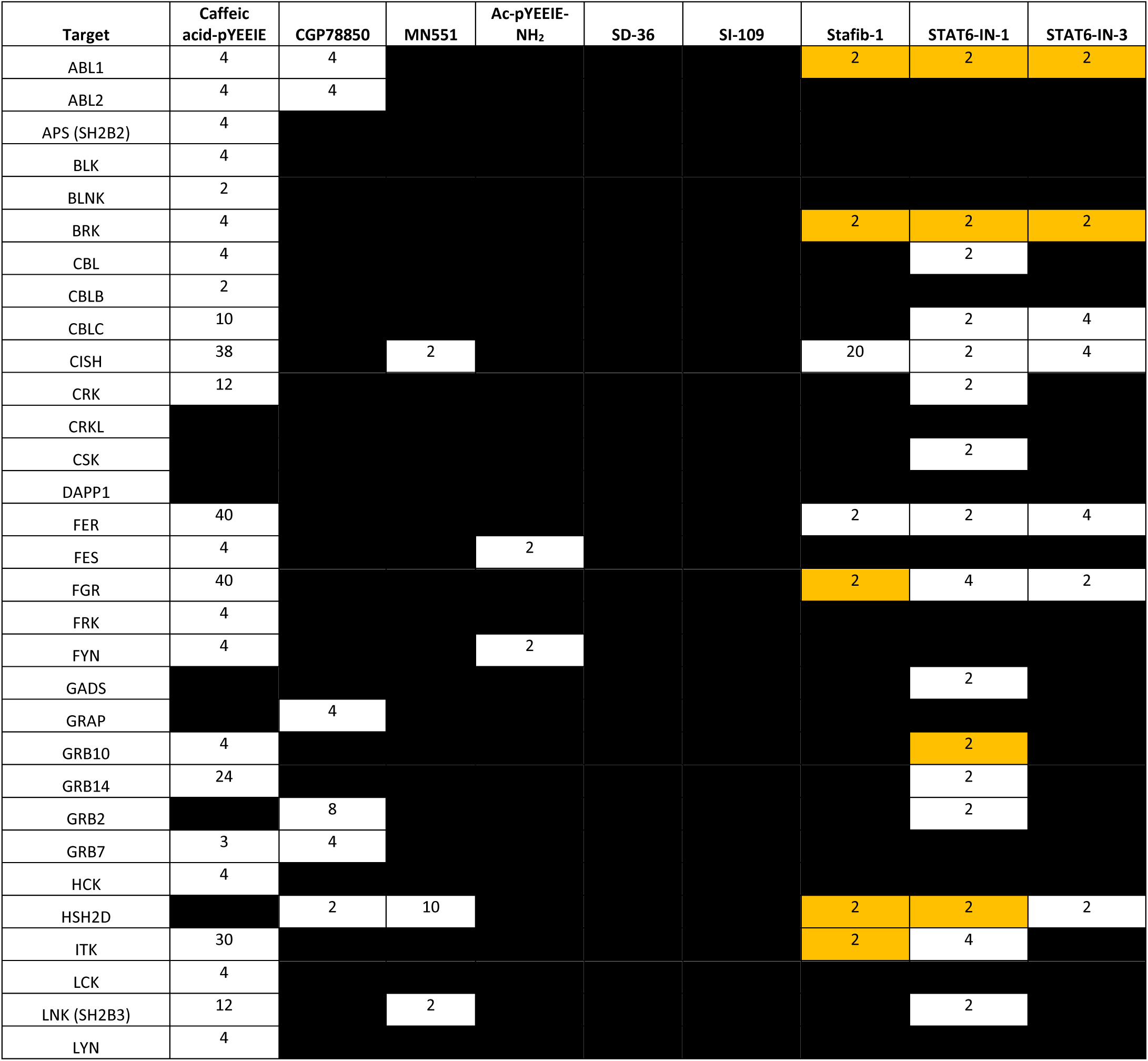
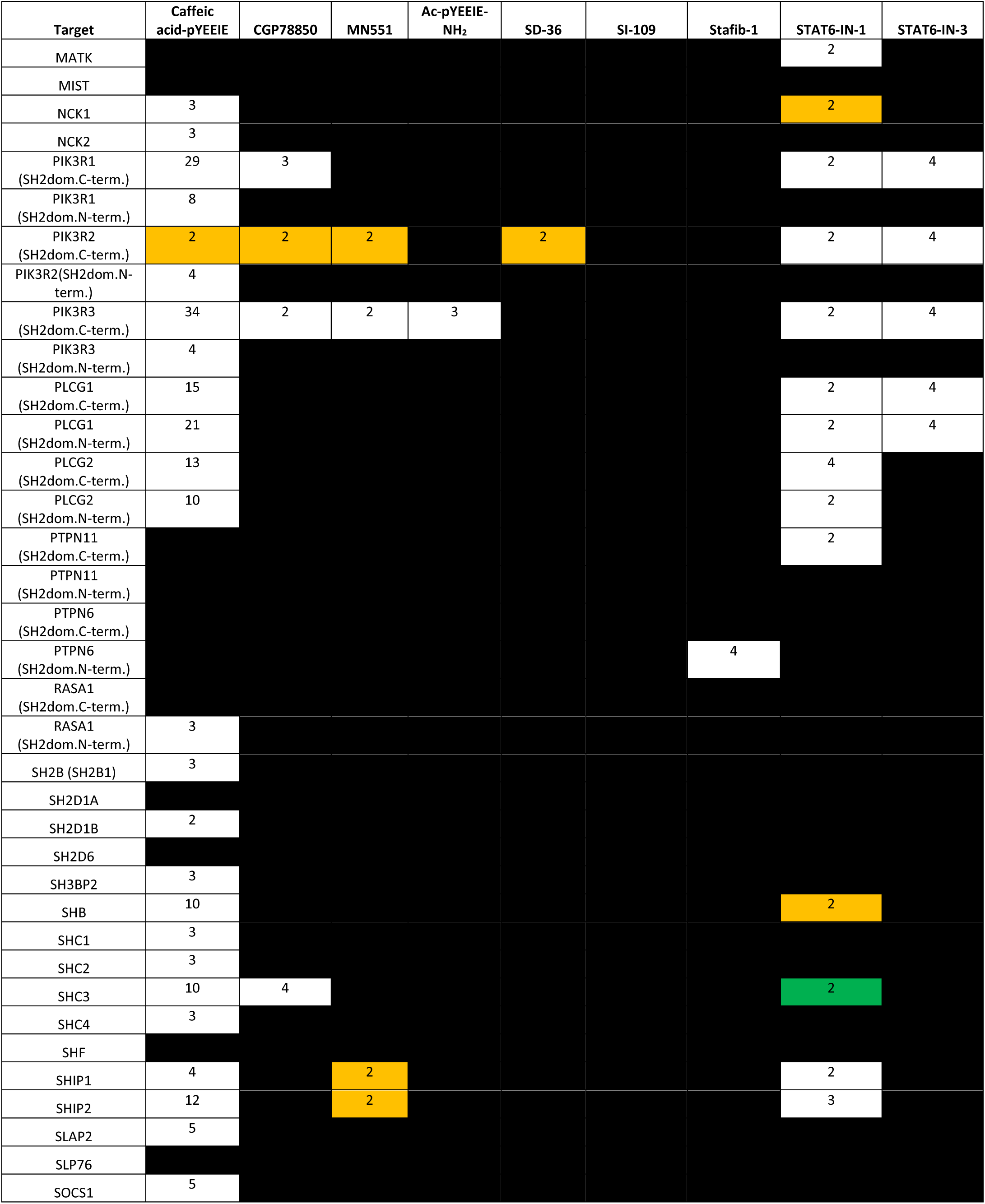
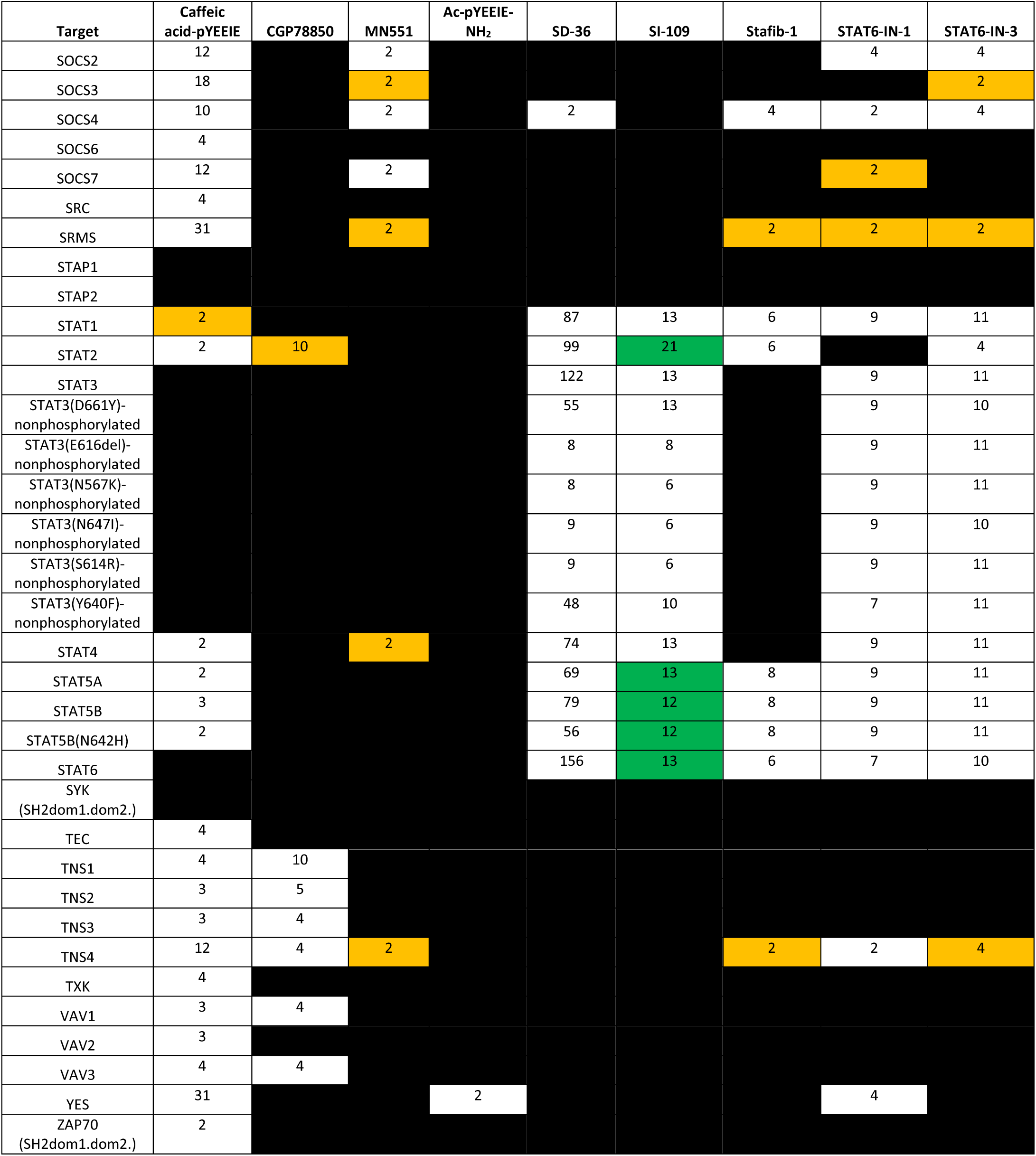
Replicate numbers for dissociation constant values for the 9 compounds tested in SH2*scan*. Independent replicate numbers (N) for each dose-response value presented in **Extended Data Table 4** are shown below and were generated from at least one independent experiment. False positives are colored in orange and false negatives are colored in green.

**Extended Data Table 6.** Literature SH2 domain binding measurements for compounds relevant to this study. Literature values for the SH2 domain binding of compounds tested or related to those tested in this study are shown below in nM. Units reported for these measurements are listed in brackets following the value in the table.

| Target | Caffeic acid-pYEEIE | CGP78850 | MN551 | Ac-pYEEIE | SD-36 | SI-109 | Stafib-1 | STAT6-IN-1 | STAT6-IN-3 |
| --- | --- | --- | --- | --- | --- | --- | --- | --- | --- |
| FYN | 84 (IC<sub>50</sub>) | >50000 (IC<sub>50</sub>) |  | 1100 (IC<sub>50</sub>) |  |  |  |  |  |
| GRB2 |  | 40 (IC<sub>50</sub>) |  |  |  |  |  |  |  |
| LCK | 42 (IC<sub>50</sub>) |  |  | 1300 (IC<sub>50</sub>) |  |  |  |  |  |
| PIK3R1 (SH2dom.N-term.) |  | >50000 (IC<sub>50</sub>) |  |  |  |  |  |  |  |
| PTPN11 (SH2dom.C-term.) |  | >50000 (IC<sub>50</sub>) |  |  |  |  |  |  |  |
| PTPN11 (SH2dom.N-term.) |  | >50000 (IC<sub>50</sub>) |  |  |  |  |  |  |  |
| SOCS2 |  |  | 2180 (K<sub>D</sub>) |  |  |  |  |  |  |
| SRC | 64 (IC<sub>50</sub>) |  |  | 2000 (IC<sub>50</sub>) |  |  |  |  |  |
| STAT1 |  |  |  |  | 1086 (K<sub>D</sub>) | 1270 (K<sub>D</sub>) |  |  |  |
| STAT2 |  |  |  |  | 4883 (K<sub>D</sub>) | 6630 (K<sub>D</sub>) |  |  |  |
| STAT3 |  |  |  |  | 44.4 (K<sub>D</sub>) | 51.2 (K<sub>D</sub>) |  |  |  |
| STAT4 |  |  |  |  | 804 (K<sub>D</sub>) | 1980 (K<sub>D</sub>) | 3290 (K<sub>i</sub>) |  |  |
| STAT5A |  |  |  |  | 6180 (K<sub>D</sub>) |  | 2420 (K<sub>i</sub>) |  |  |
| STAT5B |  |  |  |  | 11700 (K<sub>D</sub>) |  | 44 (K<sub>i</sub>) |  |  |
| STAT6 |  |  |  |  | 11325 (K<sub>D</sub>) | 3370 (K<sub>D</sub>) | 3470 (K<sub>i</sub>) | 28 (IC<sub>50</sub>) | 44 (IC<sub>50</sub>) |
| References | Park *et al.*, Bioorganic & Medicinal Chemistry Letters, 2002 | Gay *et al.*, International Journal of Cancer, 1999 | Ramachandran *et al.*, Nature Communications, 2023 | Park *et al.*, Bioorganic & Medicinal Chemistry Letters, 2002 | Bai *et al.*, Cancer Cell, 2019 | Bai *et al.*, Cancer Cell, 2019 | Elumalai *et al.*, Angewandte Chemie International Edition, 2015 | Mandal *et al.*, Journal of Medicinal Chemistry, 2015 | Mandal *et al.*, Journal of Medicinal Chemistry, 2015 |

